# Differential Impacts of Dietary Modification on Individual Metabolic Phenotypes and their Relationship to Blood Pressure: Evidence of Latent Dietary Responder Sub-phenotypes

**DOI:** 10.1101/193359

**Authors:** Ruey Leng Loo, Xin Zou, Lawrence J Appel, Jeremy K. Nicholson, Elaine Holmes

## Abstract

**Background:** Hypertension is a worldwide public health issue with significant comorbidity and mortality. We aimed to identify urinary metabolic phenotypes associated with three healthy diets and to establish their relationship to blood pressure (BP).

**Methods and Results:** —24-h urine samples from 158 participants, with pre-hypertension and hypertension, consumed a carbohydrate-rich, a protein-rich and a monounsaturated fat-rich healthy diet (6-week per diet) in randomized order, were analyzed by nuclear magnetic resonance spectroscopy. Combinations of metabolites significantly associated with each diet were identified, and associations between these metabolites and cardiovascular disease risk were established. We found coherent responses to all three diets including increased excretion of metabolites originating from vegetables/fruits, protein, tryptophan metabolism and gut microbial-mammalian co-metabolism. Proline betaine (marker of citrus fruit) was significantly inversely associated with systolic BP; 4-cresyl sulfate (gut microbial metabolite) inversely correlated with both systolic and diastolic BP; and hippurate (gut microbial metabolite) - directly associated with reduced systolic BP.

**Conclusions:** Variation in metabolic phenotypes in response to specific diets may hold clues as to the mechanisms underlying inter-individual differences in dietary response. Stratification of individuals based on diet-specific urinary phenotypes highlights the feasibility for individualized approaches to dietary therapy for lowering BP.

**Clinical Trial Registration:** This intervention study is registered at http://www.clinicaltrials.gov as NCT00051350

## INTRODUCTION

The pathogenesis of cardiovascular diseases (CVD) is multifactorial and complex. Worldwide, CVD contributes to almost a third (17 million) of the total deaths per year, and the complications of hypertension alone are accountable for over 50% of the CVD-related deaths^1^. The high global prevalence of hypertension is a major public health issue that requires urgent attention. Prevention of hypertension-related disease using dietary solutions, without the need for pharmacological intervention, is particularly desirable with regard to social and financial management of the population burden of CVD. A number of healthy diets such as the Dietary Approaches to Stop Hypertension (DASH)^2^, Mediterranean^3^ and Optimal Macronutrient Intake Trial to Prevent Heart Disease (OmniHeart)^4^ diets, have been shown to reduce blood pressure (BP); optimize serum lipid profiles; and improve cardiovascular risk scores.

Despite the fact that healthy diets have unequivocally been shown to impact beneficially on health at a population level, at the inter-individual level, the evidence is conflicting^5^. Investigation of gene-environmental interactions by metabolic phenotyping is an attractive approach as it enables better health outcomes to be achieved using stratification of individuals’ disease risks and biological phenotypes^6^. Metabolic phenotyping studies have proven valuable in investigating changes in metabolic phenotypes in response to food intake^7^, evaluating dietary modulation in gastrointestinal cancer risk^8^ and in metabolically characterizing BP^9, 10^ and obesity^11^. In an attempt to understand this variability in dietary phenotypes, we used proton nuclear magnetic resonance (^1^H NMR) spectroscopy to assess the 24-h urinary phenotype of 158 individuals in response to three different healthy diets, each consumed for a 6-week period, in a randomized cross-over feeding study. We hypothesized that comparison of healthy diets with a typical American diet or between different healthy diets would result in specific changes in the urinary metabolic phenotypes. We further hypothesize that these specific urinary metabolic phenotypes will be associated with CVD risk factors.

## METHODS

The OmniHeart Study (N=164) was a randomized, controlled, cross-over feeding study that assessed the effects of three healthy diets on the BP and lipid profiles of participants. All three OmniHeart diets had a similar nutrient composition to the established healthy DASH diet but varied in macronutrient composition. The Omniheart carbohydrate-rich diet (OmniCarb diet) provided 58% kcals from carbohydrate, 15% from protein and 27% from fat; the remaining two diets, replaced 10% of calories from carbohydrate with either protein, predominantly obtained from vegetable sources (OmniProt diet), or unsaturated fats, predominantly derived from monounsaturated fat (OmniFat diet).

A total of 158 healthy men and women, aged between 30 to 80 years, with pre-hypertension or stage 1 hypertension and without diabetes or prior CVD completed all three dietary interventions. Four 24-h urine collections were obtained, one during a screening visit (baseline) and one at the end of each dietary intervention. During the last 10 days of each dietary intervention, BP measurements on 5 days and a fasting blood specimen for full lipid profiles were also obtained. Compliance to diets was considered to be >95% for all three diets, based on self-reported data and objective data, i.e. mean levels of urinary urea nitrogen excretion, which was the highest for the OmniProt diet^4^. Baseline socio-demographic, anthropometric characteristics together with the mean changes from baseline to each dietary intervention are shown in Supplementary Table 1. All participants were recruited from the Baltimore and Boston areas, and institutional ethics committee approval was obtained for each site; all participants provided written informed consent.

### Data processing and analysis

Urine specimens were prepared and analyzed by 600 MHz ^1^H NMR spectroscopy following standard protocol^12^. The processed spectroscopic data were normalized and modelled using orthogonal partial least squares-discriminant analysis (OPLSDA). Due to the nature of cross-over study design, we then employed multilevel OPLSDA (mOPLSDA)^13^, which incorporates the variation between and within participants in the dataset to optimize visualization of dietary response. Because of the heterogeneity in clinical responses amongst the participants, a variant of statistical correlation spectroscopy^14^ known as the Statistical HOmogeneous Cluster SpectroscopY (SHOCSY)^15^ method was applied to each mOPLSDA comparison to enable the characterization of homogeneous subgroups of spectra from participants showing uniform responses (homogeneous subgroups) and to differentiate these from spectra indicating a non-uniform response (heterogeneous/variable subgroups) to each diet. Using this strategy, we compared urinary spectral data obtained at the end of each dietary intervention to baseline and between the three OmniHeart diets in a pairwise fashion and modelled these data separately for the homogeneous and the variable subsets (total of 12 models). The robustness of each model was assessed by seven-fold cross validation Q^2^ ^16^ and permutation testing (100 repetitions).

Discriminatory metabolites were extracted from each mOPLS-DA model on the basis of assessed stability of the NMR variable where at least three consecutive NMR data points showed a loading regression coefficient (r^2^) >0.3 with a significant P value of <1.85 x 10^-6^ after Sidák correction^15^. The identification of discriminatory metabolites was confirmed by spiking with authentic standard compounds, where available. Additional acquisition of a series of two-dimensional NMR experiments and a tandem mass spectrometry (MS/MS) spectra were used in conjunction with statistical correlation analysis and published databases^17^.

Since we found no significant difference in the excretion of creatinine between different OmniHeart diets and the typical American diet (P >0.5), the difference in absolute excretion of each discriminatory metabolite between each comparison was adjusted to the corresponding 24-h urinary creatinine excretion (in mmol/L). The association between the changes in discriminatory metabolites and CVD risk was established using Spearman rank correlation for homogeneous responders. In addition, the known covariate for hypertension including urinary sodium, potassium, calcium and phosphate, was also established for homogeneous and variable responders. The statistical significance of Spearman rank correlation was adjusted by Bonferroni correction (0.05/number of comparisons) to account for multiple testing. All analyses were performed using in-house software written in Matlab (version 2012a, MathWorks, Natick, MA). Full details of the NMR experimental parameters for data processing and multivariate data analysis are provided in Supplementary Information.

## RESULTS

### OmniHeart diets are associated with distinctive metabolic phenotypes compared to the typical American diet

Initially, the spectral data were modelled using standard OPLSDA in a pairwise fashion but the differentiation in the urinary spectral data attributable to diet were not captured successfully by this modelling method, most likely due to the relatively subtle changes in the OmniHeart diets when placed in context of the magnitude of overall variation introduced by the diversity of genetic and environmental exposures. Application of SHOCSY to identify inherent subgroups in each dietary class, aiming to distinguish those individuals with a similar response (homogeneous subgroup) to a specific diet, indicated that the majority of the participants showed a coherent metabolic response to all of the healthy OmniHeart diets, with the response to the OmniProt diet being the most variable between participants when compared with the baseline profile: 71.5% (N=113) for OmniProt, 80.4% (N=127) for OmniFat and 86.7% homogeneous response (N=137) for OmniCarb. The remaining individuals who were characterized by metabolic phenotypes that differed from these homogeneous subgroups were denoted as ‘variable’ subgroups/individuals, i.e. 45 for OmniProt, 31 for OmniFat, and 21 for OmniCarb.

We firstly examined the urinary metabolite profiles of the homogeneous subgroups obtained after following each of the OmniHeart diets and compared them to their corresponding baseline metabolic phenotypes using mOPSLDA. The mOPLSDA models for all three diets were highly significant based on permutation testing (P<10^-5^ for all homogeneous subgroups) and the discriminatory metabolites characteristic for each of the diets were identified (Supplementary Table 2). The origin of the discriminatory metabolites for these three healthy diets over the metabolic profiles at baseline could be predominantly assigned to i) diet: including proline betaine (PB), *N*-acetyl-S-methyl-L-cysteine sulfoxide (NAMCO), S-methyl-L-cysteine-S-oxide (MCO), creatine and carnitine; ii) tryptophan-nicotinamide-adenine dinucleotide (NAD) degradation e.g. *N*-methyl-2-pyridone-5-carboxamide (M2PY), *N*-methyl nicotinic acid (NMNA), and *N*-methyl nicotinamide (NMND); and iii) gut microbial-mammalian metabolism e.g. hippurate, 4-hydroxyphenylacetic acid (HPA) and dimethylglycine (DMG). PB was the only metabolite uniformly increased in the urinary profiles for all three OmniHeart homogeneous subgroups, which is consistent with increased citrus fruit consumption^18^ in all three diets. Increased excretion of hippurate, PB, MCO, and NMNA; and reduced excretion of HPA, M2PY, *N*-acetylglycoprotein, (NAC), NMND and an unknown metabolite (U1), consisting of three singlets δ0.90, δ0.94 and δ0.95 (as identified using statistical correlation), were observed in the homogeneous subgroups of OmniCarb diet when compared to the baseline. Each OmniHeart diet was unique in its biochemical consequences; the homogeneous subgroup of OmniProt was characterized by increased excretion of carnitine, creatine, DMG, NAMCO with reduced excretion of U1, whereas OmniFat showed increased excretion of DMG but reduced excretion of U1 when compared to the baseline profile.

**Table 1:**
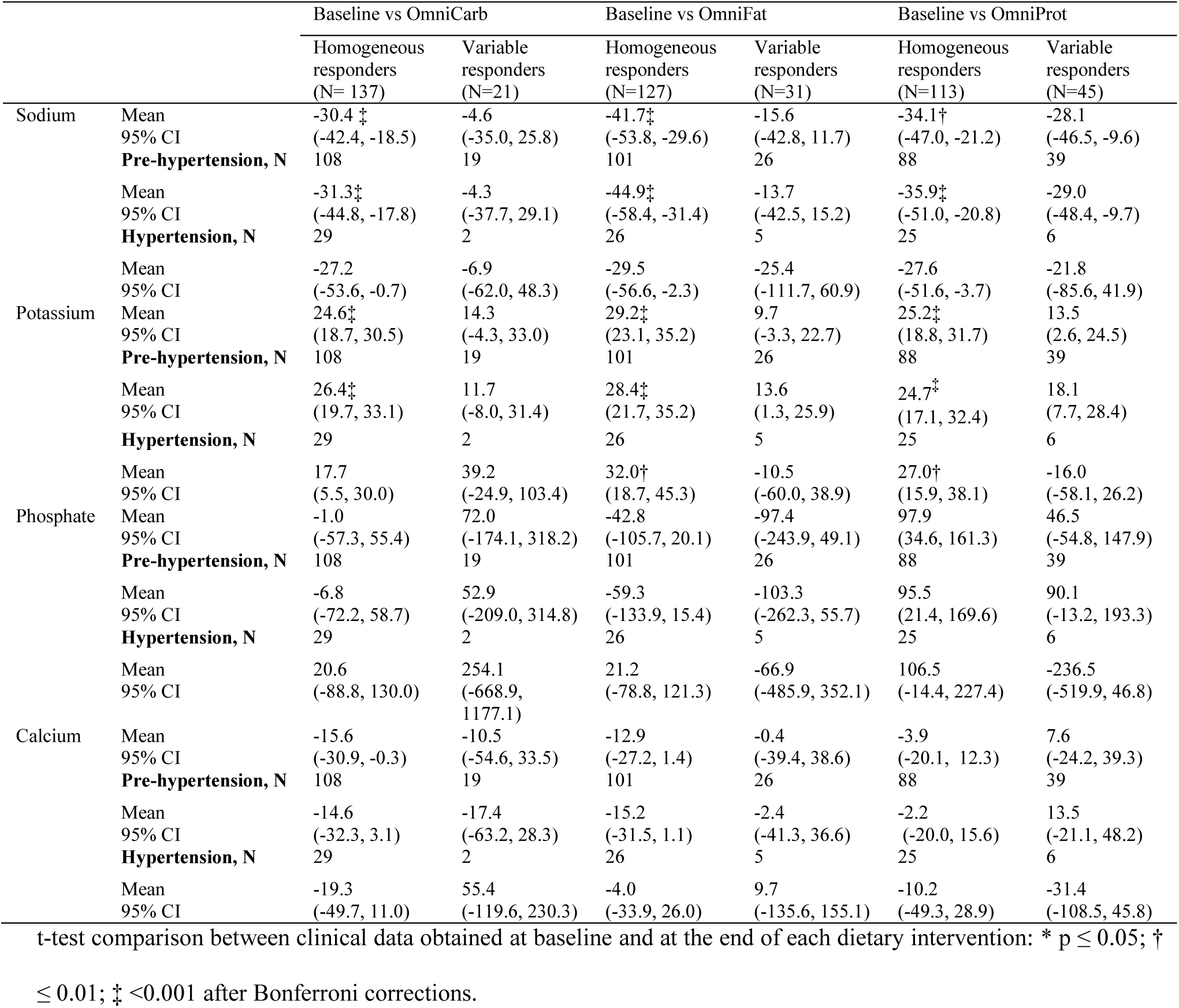
Mean changes in urinary electrolyte levels for sodium, potassium, phosphate and calcium (95% confident interval) for homogeneous and variable subgroups for each of the specific OmniHeart diets.

**Table 2:**
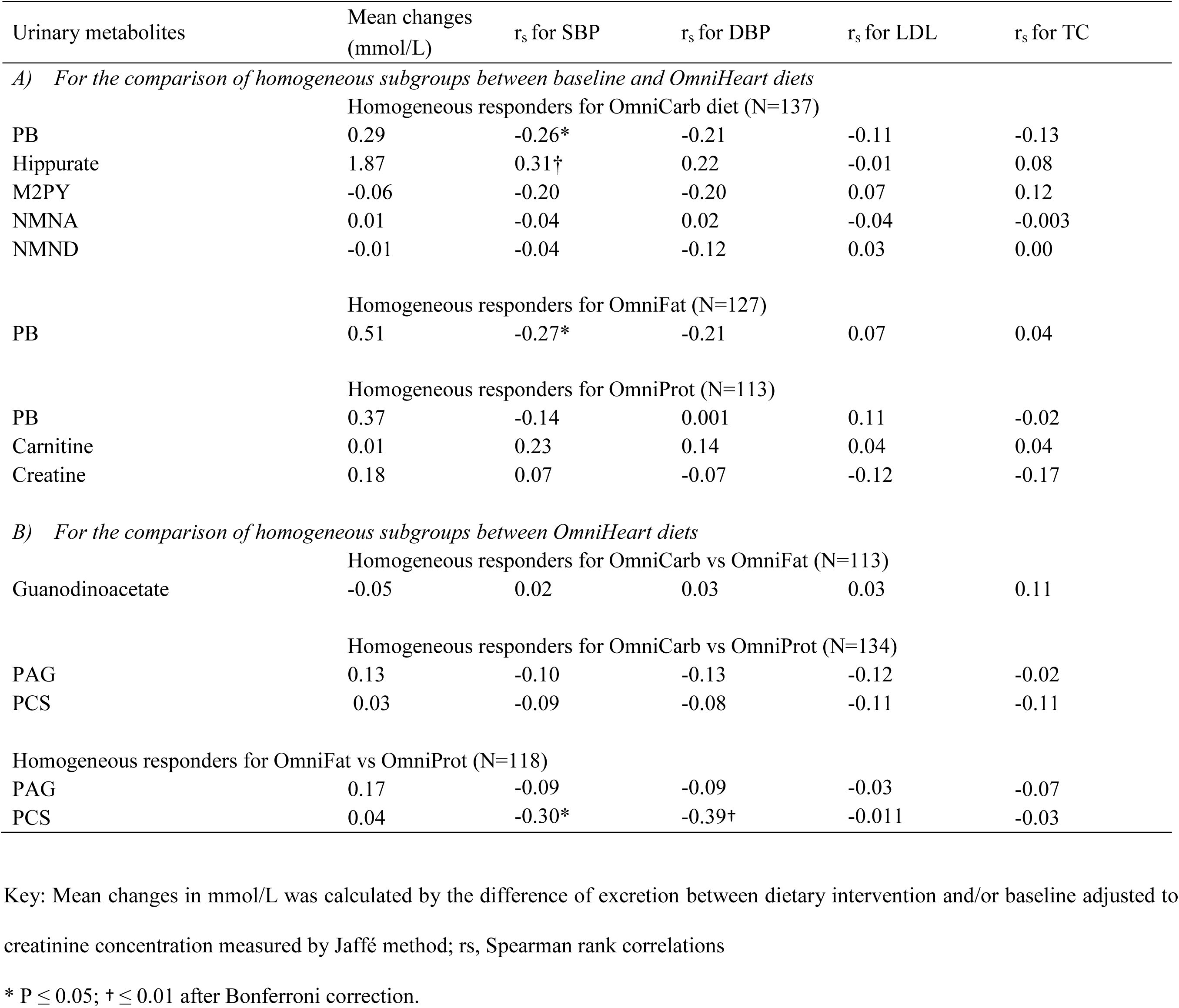
Association between urinary discriminatory metabolites and CVD risk factors by Spearman rank correlation for A) comparison between baseline and OmniHeart diets; B) comparison between different OmniHeart diets.

| Urinary metabolites | Mean changes<br>(mmol/L) | r<sub>s</sub> for SBP | r<sub>s</sub> for DBP | r<sub>s</sub> for LDL | r<sub>s</sub> for TC |
| --- | --- | --- | --- | --- | --- |
| *A) For the comparison of homogeneous subgroups between baseline and OmniHeart diets* |  |  |  |  |  |
| Homogeneous responders for OmniCarb diet (N=137) |  |  |  |  |  |
| PB | 0.29 | -0.26* | -0.21 | -0.11 | -0.13 |
| Hippurate | 1.87 | 0.31† | 0.22 | -0.01 | 0.08 |
| M2PY | -0.06 | -0.20 | -0.20 | 0.07 | 0.12 |
| NMNA | 0.01 | -0.04 | 0.02 | -0.04 | -0.003 |
| NMND | -0.01 | -0.04 | -0.12 | 0.03 | 0.00 |
| Homogeneous responders for OmniFat (N=127) |  |  |  |  |  |
| PB | 0.51 | -0.27* | -0.21 | 0.07 | 0.04 |
| Homogeneous responders for OmniProt (N=113) |  |  |  |  |  |
| PB | 0.37 | -0.14 | 0.001 | 0.11 | -0.02 |
| Carnitine | 0.01 | 0.23 | 0.14 | 0.04 | 0.04 |
| Creatine | 0.18 | 0.07 | -0.07 | -0.12 | -0.17 |
| *B) For the comparison of homogeneous subgroups between OmniHeart diets* |  |  |  |  |  |
| Homogeneous responders for OmniCarb vs OmniFat (N=113) |  |  |  |  |  |
| Guanodinoacetate | -0.05 | 0.02 | 0.03 | 0.03 | 0.11 |
| Homogeneous responders for OmniCarb vs OmniProt (N=134) |  |  |  |  |  |
| PAG | 0.13 | -0.10 | -0.13 | -0.12 | -0.02 |
| PCS | 0.03 | -0.09 | -0.08 | -0.11 | -0.11 |
| Homogeneous responders for OmniFat vs OmniProt (N=118) |  |  |  |  |  |
| PAG | 0.17 | -0.09 | -0.09 | -0.03 | -0.07 |
| PCS | 0.04 | -0.30* | -0.39† | -0.011 | -0.03 |

Key: Mean changes in mmol/L was calculated by the difference of excretion between dietary intervention and/or baseline adjusted to creatinine concentration measured by Jaffé method; rs, Spearman rank correlations
\* $P \leq 0.05$ ; † $\leq 0.01$ after Bonferroni correction.

As expected, the model statistics comparing the urinary spectral data for the variable subgroups for each of the OmniHeart diets typically elicited fewer discriminatory metabolites characterizing the dietary intervention than the homogeneous subgroups. In fact, some of the discriminatory metabolites in the variable subgroups showed an inverse pattern to those found in homogeneous subgroups. For example, in the case of the OmniCarb diet comparison, the variable subgroup (OmniCarb-var) was characterized by reduced excretion of hippurate, while the homogeneous subgroup (omniCarb-homo) manifested increased excretion of hippurate (a gut-microbial-mammalian co-metabolite). Another example is the comparison of the homogeneous and variable subgroups for both the OmniFat and OmniProt diets, where the variable subgroups showed reduced excretion of PB, whilst increased PB excretion was strongly associated with the homogeneous subgroups (Supplementary Table 2).

We compared the 24-h urinary excretion of PB between the homogeneous and variable groups for each pairwise comparison. We found 15 of the 21 individuals (71.4%) from the OmniCarb-var group showed a 24-h urinary excretion of less than 95% confidence interval (CI) of the OmniCarb-homo group. A similar trend was observed for the OmniFat (21/31, 67.7%) and OmniProt (35/45, 77.8%) variable groups. This gave an estimation of individuals who were potentially non-adherent to each diet based on spectral evidence of lower fruit and polyphenol intake and an overall level of non-adherence of 9.5% to 22.2%, depending of the type of diets. We excluded those individuals who were deemed non-adherent to the diets and recalculated additional mOPLSDA models using the adherent individuals within the variable subgroups for OmniFat (N=10) and OmniProt (N=10) respectively, but not for OmniCarb-var, due to the small number of participants remaining after exclusion of non-adherent individuals. We found these variable subgroups were generally represented by different urinary phenotypes to the core homogenous subgroups; the OmniProt variable subgroup manifested higher excretion of MCO, NAMCO, and creatine but lower excretion of NAC; while the OmniFat variable subgroup was characterized by higher excretion of hippurate but lower excretion of NAC and unknown metabolite consistent with scyllo-inositol at δ3.34s when compared to the corresponding baseline variable subgroups (data not shown).

### Urinary Metabolic Variations Reflect Inter-individual Differences in Response to Clinical Benefits

We investigated the changes in CVD risk for the homogeneous and variable subgroups separately. The homogeneous groups generally showed a stronger association and a more significant reduction from baseline in CVD risk factors than the variable subgroups for: SBP (-8.5mmHg for OmniCarb; -9.7mmHg for OmniFat; and -10.0mmHg for OmniProt); DBP (-4.4mmHg for OmniCarb; -5.1mmHg for OmniFat; and -5.6mmHg for OmniProt); LDL (-12.0mg/dL for OmniCarb; -14.6mg/dL for OmniFat; and -14.9mg/dL for OmniProt); and total cholesterol, TC (-13.2mmHg for OmniCarb; -16.6mmHg for OmniFat; and -20.8mmHg for OmniProt) for all OmniHeart diets, P values all <10^-10^; and triglycerides -16.6mg/dL (P=0.002) for OmniProt diet, Figure 1. The OmniHeart diets also showed greater benefit for individuals with higher CVD risk compared to than those with lower risk, e.g. participants who were hypertensive showed a greater reduction in both SBP and DBP than those who were pre-hypertensive; whilst those with non-optimal lipid profiles showed greater reduction in both triglycerides and TC levels, Supplementary Figure 1.

**Figure 1:**
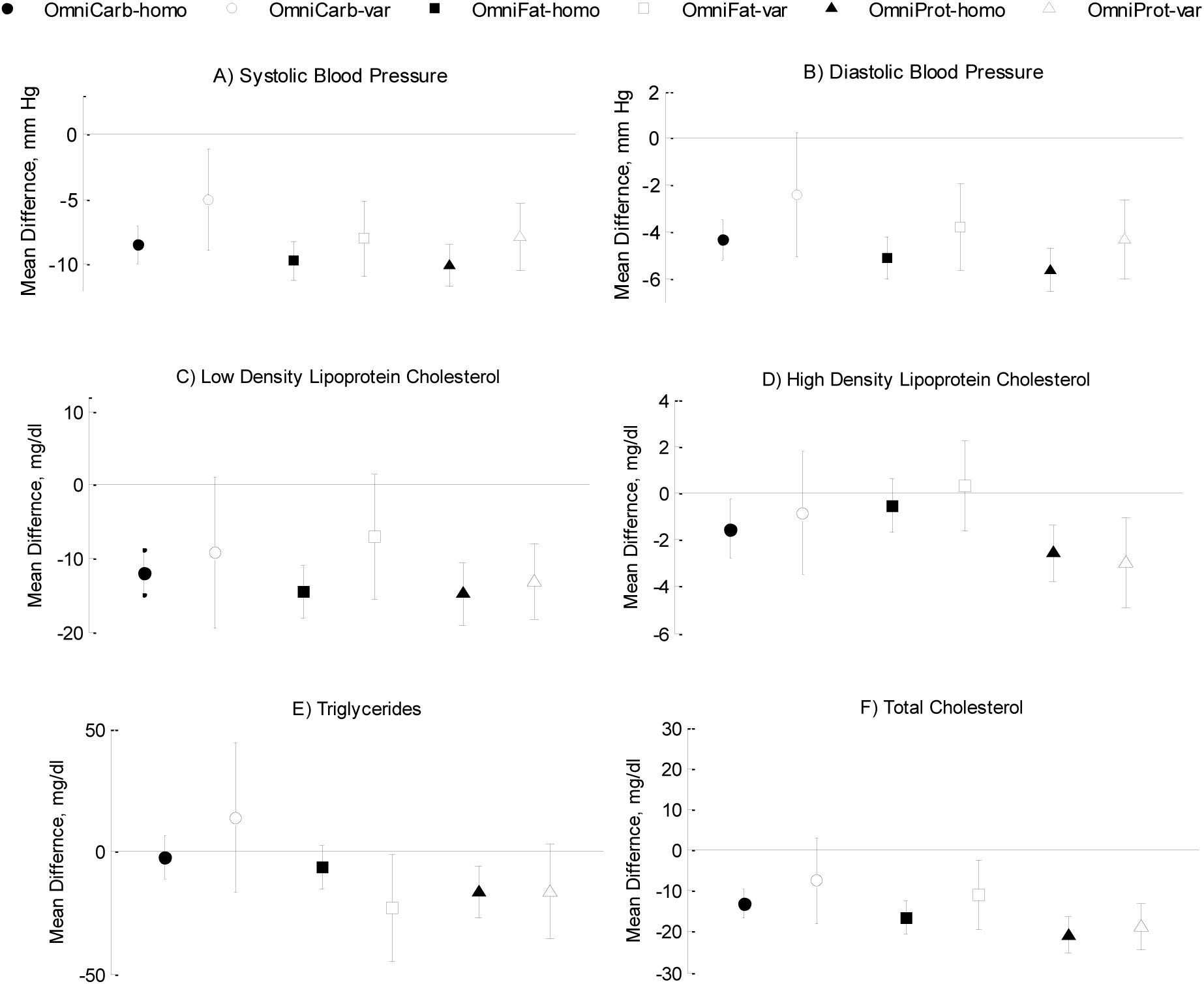
Mean changes in CVD risk factors stratified by homogeneous and variable subgroups for each OmniHeart diet measured as dietary responses in compared to baseline for: A) SBP; B) DBP; C) LDL; D) HDL; E) triglycerides; and F) TC. Key: Error bars indicate 95% confidence interval. Discrepancies in the number of participants were due to missing data in the CVD risk scores. T-test comparison was not performed when N<5 and significant P values are italicized.

For all the variable subgroups, a dampened reduction in CVD risk was observed when compared to the corresponding homogeneous diet subgroups. Nevertheless, for SBP and DBP a significant (P<10^-3^) level of reduction was observed for all variable subgroups except the OmniCarb variable subgroup. Other CVD risk factors generally showed no significant reduction for the variable subgroups when compared with the baseline. This general lack of significance of clinical benefit in the variable groups may be due to the reduced sample size, with the exception of the OmniProt variable subgroup, which demonstrated a significant lowering of LDL, HDL and TC; and for the OmniFat variable subgroup, which showed a significant reduction in the response of triglycerides and TC.

We also investigated urinary electrolytes known to have an effect on BP. For each OmniHeart diet, we observed significant reduction in urinary sodium and a significant increase in urinary potassium among the homogeneous subgroups, Table 1. The overall mean changes of urinary sodium and potassium were only significant amongst participants who were hypertensive and showed homogeneous responses to diets. These changes were insignificant for all other subgroups. Trends toward reduced sodium and increased potassium excretion were observed in the variable subgroups, but these changes were typically not significant. Changes in the urinary calcium and phosphate were not statistically significant for any dietary group.

We found that individuals who were consistently classed as a homogeneous responder to all three OmniHeart diets (60.1%, N=95) generally manifested a greater reduction in SBP, DBP, LDL and TC levels than those that showed homogeneous responses to just one or two OmniHeart diets (Supplementary Figure 2). Similar to the results observed earlier among the homogeneous subgroups to specific OmniHeart diets, the reduction in CVD risk was generally greater in those high risk subgroups e.g. those who were hypertensive or with a non-optimal lipid profile, indicating the potential benefit of these three healthy diets. For the variable subgroup, after excluding spectral data with low urinary excretion of PB, indicating non-adherent, we found 23(14.5%) individuals showed variable responses to one or more OmniHeart diet.

### Distinct changes in urinary metabolic phenotype were observed between OmniHeart diets

Pairwise comparison between OmniHeart diets showed consistent differences in diet-associated metabolic phenotypes (P<10^-5^). The homogeneous subgroup of OmniProt diet was generally characterized by higher excretion of urinary creatine (indicative of protein intake); M2PY and two gut microbial mammalian co-metabolites, phenylacetylglutamine (PAG) and 4-cresyl sulfate (PCS); whilst the homogeneous subgroup of OmniCarb consistently showed higher excretion of hippurate and guanodinoacetate when compared to other OmniHeart homogeneous subgroups, Supplementary Table 3 and 4. As expected, the differences in the markers for dietary intake of cruciferous vegetables, MCO and NAMCO, and markers for citrus fruit intake, PB, observed when comparing urine of OmniHeart diets with the baseline profiles, were generally not observed for pairwise comparisons between the OmniHeart diets, since all three OmniHeart diets contained more fruits and polyphenols than the baseline diet.

### Correlations of 24h Urinary Excretion of Discriminatory Metabolites with CVD Risk Factors

We then quantified the 24-h urinary excretion of the 8 key discriminatory metabolites using the NMR signal and assessed whether there was a direct relationship between the urinary metabolite characteristics for different OmniHeart diets and their reduction in CVD risk factors. We found a significant inverse association between the PB excretion and SBP for both the OmniCarb and OmniFat diets when compared to baseline, Table 2. Although a similar trend was observed for the OmniProt diet, this association was not statistically significant. We also found a significant direct association between the changes in the excretion of hippurate with the reduction in SBP for OmniCarb diet when compared to baseline. Although there was a trend in the association between M2PY and the reduction in DBP for OmniCarb diet; and excretion of carnitine with the changes in SBP for OmniProt diet, when compared to baseline, these trends were not significant after correction for multiple testing. No significant associations were observed between the changes in these urinary metabolites and changes in the LDL and TC. Analyses were not performed for HDL and triglycerides as our results showed the changes in these clinical parameters were generally not significant among the homogeneous subgroups for any of the three OmniHeart diets compared to baseline. Significant inverse associations were observed for changes in the excretion of PCS and the reduction in SBP and DBP for the pairwise comparison between OmniProt and OmniFat.

## DISCUSSION

Poor diet and other lifestyle factors contribute to the high burden of hypertension and elevated CVD risk. Dietary strategies to improve risk factors are desirable and have led to the development of healthy diets, such as the DASH diet, which is rich in fruits, vegetables and low-fat dairy products and reduced in saturated fat and cholesterol. However, despite the effectiveness of these diets at a population level, there is substantial inter-individual variability. In this study, we demonstrate a method for stratification of individuals based on diet-specific urinary phenotypes in order to better characterize CVD risk. Using a statistical correlation algorithms, SHOCSY which incorporates stringent Sidák correction to minimise false positive results and identifies a core group of ‘homogeneous’ responders, we have shown remarkably consistent modulatory effects on the urinary metabolome when individuals adopted healthy dietary patterns. The homogeneous group represented the majority of the participants for each of the healthy OmniHeart diets with a minority group of ‘variable’ responders. We show that homogeneous group for each diet typically demonstrated stronger correlations between the metabolic response to a healthy diet and improvement in clinical measurements. We identified generic ‘markers’ for healthy diet deriving directly from the diet itself such as MCO and NAMCO from cruciferous vegetables^19^ and PB from citrus fruits^18^ in the homogeneous OmniHeart diets when compared the typical American diet. When pairwise responses between the different OmniHeart diets were evaluated, these markers were no longer discriminatory since all OmniHeart diets involved increased fruit and vegetable intake when compared to a typical American diet. As expected, higher urinary levels of carnitine and creatine, both known indices of increased protein intake^20^, remained discriminatory in the comparison of the OmniProt and baseline diets as well as the comparisions of OmniProt to OmniCarb and/or OmniFat. The high protein intake of the OmniProt diet (10% kcal higher than the other diets) was likely responsible.

We also observed differences in metabolites related to the tryptophan-NAD pathway: reduced excretion of NMND and M2PY but increased excretion of NMNA among the homogeneous responders to the OmniCarb diet compared to the baseline. Bartus *et al* found that the ingestion of 1-methylnicotinamide in hypertriglyceridemic rats resulted in an increase of 1-methylnicotinamide and its metabolites such as M2PY, concomitant with a significant reduction in serum triglycerides. They also found that ingestion of 1-methylnicotinamide in both diabetic and hypertriglyceridemic rats ameliorated the nitric oxide dependent vasodilation, a key surrogate marker for atherosclerosis and improved CVD health^21^. Increased urinary excretion M2PY and NMND with a decreased in NMNA has also been observed in type 2 diabetes patients^22^. In contrast, we, and others, did not find such a correlation between urinary excretion and serum lipid profiles^23^.

We observed that the participants in the variable subgroups demonstrated some metabolites showing an opposing metabolite pattern to the homogeneous groups. These individuals typically showed a dampened clinical response to the dietary interventions. The reasons for variable response to dietary intervention was obvious for some participants, for example, reduced PB excretion appeared to indicate a lack of dietary adherence with respect to citrus fruit intake, as PB is excreted mainly unchanged with the production of some minor metabolites in the urine^24,25^. We therefore used PB excretion as a proxy marker for non-adherence to the OmniHeart diets and found that the majority of the participants in the variable response groups showed a reduction in 24-h urinary excretion of PB when compared to the homogeneous groups. These individuals corresponded to an overall estimated non-adherence of 9.5% (n=15), 13.3% (n=21) and 22.2% (n=35) for OmniCarb, OmniFat and OmniProt diets, respectively. These values were considerably higher than the <5% non-adherence estimated from the self-reported data^4^. Based on the urinary metabolic phenotypes, identification of individuals showing limited clinical responses due to non-adherence to dietary intervention is therefore made possible. This offers the feasibility of a rational stratification strategy for clinicians to apply an alternative clinical intervention for optimizing CVD risk. We also observed that a small proportion of the individuals in the variable groups who appeared to be adherent to diet, based on the measured PB excretion, remained discordant from the homogeneous subgroups in terms of their metabolic phenotype. These subgroups, albeit small in proportion, may indicate individuals who differ in their metabolism of dietary components from the majority of the population. In these subgroups, we observed diet-specific differences such as hippurate, that exhibited contrasting behaviour in the homogenous and variable groups, and which may be indicative of inter-individual differences in gut-microbial related metabolic activity. We also found the homogenous OmniProt group generally showed an increased excretion of PAG and PCS when compared to other OmniHeart diets but not the typical American diet. The differences in gut-microbial co-metabolites may indicate the differences in putrefaction of protein following an increase in protein content of the OmniProt diet^26^. Numerous studies have reported associations between gut bacteria and obesity, diabetes and BP. Higher urinary excretion of hippurate has generally been associated with lean phenotypes in both animals^27^ and humans^11^. A large-scale cross sectional study also reported an inverse association between excretion of hippurate and BP^9^. This is consistent with our observation of increased hippurate excretion following adherence to the OmniCarb diet and its direct association with a reduction in SBP.

Large-scale genome wide association studies have widely demonstrated genetic variation in response to drug treatments, particularly in the areas of anti-cancer^28^ and psychotropic drugs^29^. Similarly, variation in response to diets has also been observed based on genetic studies^30^. These variations in response are mostly attributed to genetic polymorphisms. However, to the best of our knowledge, our study is the first study that shows non-genetic variation in the urinary metabolic phenotype in response to healthy diets. Currently, nutritional recommendations are based on providing an average nutrient requirement and are typically aimed at prevention of chronic diseases and/or maintenance of healthy lifestyle of the general population. However, evidence from our results clearly indicates that approximately 14.5% of the participants did not respond in the ‘typical’ manner to one or more of the healthy OmniHeart diets despite being adherent to the diets. Extending this concept, one can envisage that further characterization of inter-individual response to healthy diet as determined by individual’s phenotypic patterns and further determining an individual’s longitudinal phenotypic stability prior to a dietary healthy intervention would need to be developed for the identification of latent sub-phenotypes. This may confer a public health benefit that has the potential to provide a personalized approach to dietary recommendations aimed at optimizing prevention of CVD and related disorders.

While our current study discovered phenotypes associated with BP and healthy dietary patterns, the mechanistic connection between these metabolites and BP regulation require further elucidation. Since by design, the weight of participants remained the same throughout the study, models were not adjusted for body mass index. We also did not adjust for socioeconomic status based on a previous large scale cross sectional study, where years of education were used as a proxy for socioeconomic status, demonstrated that the inverse association with BP was explained mostly by dietary differences^31^. The inclusion of individuals from a high CVD risk group such as African American (~50%) and pre-hypertensive patients (~80%) add strength to the general applicability of our stratification strategy, although we recognize that our findings have not been validated using an independent dataset. Nevertheless, provision of all meals to participants, inclusion of 24-h urine collection, and the randomized cross-over design all add rigor to the study, which represents one of the largest dietary interventions of its kind.

### Conclusions

The application of metabolic phenotyping to a landmark dietary intervention study, coupled with statistical stratification of the patient cohorts enabled the identification of individuals who were homogeneous in response to healthy diets and provided a tool for identifying individuals who exhibited a non-uniform response to the diets. While the majority of individuals (>60%) showed a homogeneous response to all OmniHeart diets, a small proportion of individuals manifested variable dietary responses despite being adherent to the diet. We propose that this difference could be partially attributable to differential gut microbial activities based on the urinary metabolite profiles. Using this NMR-based urinary metabolic phenotyping strategy, differentiation in the excretion of urinary metabolites enabled the assessment of CVD risk factors such as BP. In conclusion, variation in metabolic phenotypes in response to specific diets may hold clues as to the mechanisms underlying inter-individual differences in dietary response and provide a clinically actionable framework to develop tailored dietary interventions designed to reduce BP and CVD risk.

## ACKNOWLEDGEMENTS

We thank Miss T. Yap for her contribution to the sample preparation for NMR analyses. This manuscript was prepared using OmniHeart research materials obtained from the NHLBI Biologic Specimens and Data Repository Information Coordinating Centre and does not necessarily reflect the opinions or views of the NHLBI.

## REFERENCES

1. WHO. A global brief on hypertension. Silent killer, global public health crisis. 2013; WHO/DCO/WHD/2013.2

2. Appel LJ, Moore TJ, Obarzanek E, Vollmer WM, Svetkey LP, Sacks FM, Bray GA, Vogt TM, Cutler JA, Windhauser MM, Lin PH, Karanja N. A clinical trial of the effects of dietary patterns on blood pressure. Dash collaborative research group. N Engl J Med. 1997;336:1117–1124

3. Grosso G, Mistretta A, Frigiola A, Gruttadauria S, Biondi A, Basile F, Vitaglione P, D'Orazio N, Galvano F. Mediterranean diet and cardiovascular risk factors: A systematic review. Crit Rev Food Sci Nutr. 2014;54:593–610

4. Appel LJ, Sacks FM, Carey VJ, Obarzanek E, Swain JF, Miller ER, 3rd, Conlin PR, Erlinger TP, Rosner BA, Laranjo NM, Charleston J, McCarron P, Bishop LM. Effects of protein, monounsaturated fat, and carbohydrate intake on blood pressure and serum lipids: Results of the omniheart randomized trial. Jama. 2005;294:2455–2464

5. Masson LF, McNeill G, Avenell A. Genetic variation and the lipid response to dietary intervention: A systematic review. Am J Clin Nutr. 2003;77:1098–1111

6. Nicholson JK. Global systems biology, personalized medicine and molecular epidemiology. Mol Syst Biol. 2006;2:52

7. Heinzmann SS, Merrifield CA, Rezzi S, Kochhar S, Lindon JC, Holmes E, Nicholson JK. Stability and robustness of human metabolic phenotypes in response to sequential food challenges. J Proteome Res. 2012;11:643–655

8. O'Keefe SJ, Li JV, Lahti L, Ou J, Carbonero F, Mohammed K, Posma JM, Kinross J, Wahl E, Ruder E, Vipperla K, Naidoo V, Mtshali L, Tims S, Puylaert PG, DeLany J, Krasinskas A, Benefiel AC, Kaseb HO, Newton K, Nicholson JK, de Vos WM, Gaskins HR, Zoetendal EG. Fat, fibre and cancer risk in african americans and rural africans. Nat Commun. 2015;6:6342

9. Holmes E, Loo RL, Stamler J, Bictash M, Yap IKS, Chan Q, Ebbels T, De Iorio M, Brown IJ, Veselkov KA, Daviglus ML, Kesteloot H, Ueshima H, Zhao L, Nicholson JK, Elliott P. Human metabolic phenotype diversity and its association with diet and blood pressure. Nature. 2008;453:396–400

10. Menni C, Graham D, Kastenmuller G, Alharbi NH, Alsanosi SM, McBride M, Mangino M, Titcombe P, Shin SY, Psatha M, Geisendorfer T, Huber A, Peters A, Wang-Sattler R, Xu T, Brosnan MJ, Trimmer J, Reichel C, Mohney RP, Soranzo N, Edwards MH, Cooper C, Church AC, Suhre K, Gieger C, Dominiczak AF, Spector TD, Padmanabhan S, Valdes AM. Metabolomic identification of a novel pathway of blood pressure regulation involving hexadecanedioate. Hypertension.2015;66:422–429

11. Elliott P, Posma JM, Chan Q, Garcia-Perez I, Wijeyesekera A, Bictash M, Ebbels TM, Ueshima H, Zhao L, van Horn L, Daviglus M, Stamler J, Holmes E, Nicholson JK. Urinary metabolic signatures of human adiposity. Sci Transl Med. 2015;7:285ra262

12. Beckonert O, Keun HC, Ebbels TM, Bundy J, Holmes E, Lindon JC, Nicholson JK. Metabolic profiling, metabolomic and metabonomic procedures for nmr spectroscopy of urine, plasma, serum and tissue extracts. Nat Protoc. 2007;2:2692–2703

13. van Velzen EJ, Westerhuis JA, van Duynhoven JP, van Dorsten FA, Hoefsloot HC, Jacobs DM, Smit S, Draijer R, Kroner CI, Smilde AK. Multilevel data analysis of a crossover designed human nutritional intervention study. J Proteome Res. 2008;7:4483–4491

14. Cloarec O, Dumas ME, Craig A, Barton RH, Trygg J, Hudson J, Blancher C, Gauguier D, Lindon JC, Holmes E, Nicholson J. Statistical total correlation spectroscopy: An exploratory approach for latent biomarker identification from metabolic ^1^h nmr data sets. Anal Chem. 2005;77:1282–1289

15. Zou X, Holmes E, Nicholson JK, Loo RL. Statistical homogeneous cluster spectroscopy (shocsy): An optimized statistical approach for clustering of (1)h nmr spectral data to reduce interference and enhance robust biomarkers selection. Anal Chem. 2014;86:5308–5315

16. Trygg J, Wold S. O2-pls, a two-block (x-y) latent variable regression (lvr) method with an integral osc filter. Journal of Chemometrics. 2003;17:53–64

17. Wishart DS, Tzur D, Knox C, Eisner R, Guo AC, Young N, Cheng D, Jewell K, Arndt D, Sawhney S, Fung C, Nikolai L, Lewis M, Coutouly MA, Forsythe I, Tang P, Shrivastava S, Jeroncic K, Stothard P, Amegbey G, Block D, Hau DD, Wagner J, Miniaci J, Clements M, Gebremedhin M, Guo N, Zhang Y, Duggan GE, Macinnis GD, Weljie AM, Dowlatabadi R, Bamforth F, Clive D, Greiner R, Li L, Marrie T, Sykes BD, Vogel HJ, Querengesser L. Hmdb: The human metabolome database. Nucleic Acids Res. 2007;35:D521–526

18. Heinzmann SS, Brown IJ, Chan Q, Bictash M, Dumas ME, Kochhar S, Stamler J, Holmes E, Elliott P, Nicholson JK. Metabolic profiling strategy for discovery of nutritional biomarkers: Proline betaine as a marker of citrus consumption. Am J Clin Nutr. 2010;92:436–443

19. Edmands WM, Beckonert OP, Stella C, Campbell A, Lake BG, Lindon JC, Holmes E, Gooderham NJ. Identification of human urinary biomarkers of cruciferous vegetable consumption by metabonomic profiling. J Proteome Res. 2011;10:4513–4521

20. Stella C, Beckwith-Hall B, Cloarec O, Holmes E, Lindon JC, Powell J, vanderOuderaa F, Bingham S, Cross AJ, Nicholson JK. Susceptibility of human metabolic phenotypes to dietary modulation. J. Proteome Res. 2006;5:2780–2788

21. Bartus M, Lomnicka M, Kostogrys RB, Kazmierczak P, Watala C, Slominska EM, Smolenski RT, Pisulewski PM, Adamus J, Gebicki J, Chlopicki S. 1-methylnicotinamide (mna) prevents endothelial dysfunction in hypertriglyceridemic and diabetic rats. Pharmacol Rep. 2008;60:127–138

22. Salek RM, Maguire ML, Bentley E, Rubtsov DV, Hough T, Cheeseman M, Nunez D, Sweatman BC, Haselden JN, Cox RD, Connor SC, Griffin JL. A metabolomic comparison of urinary changes in type 2 diabetes in mouse, rat, and human. Physiol Genomics. 2007;29:99–108

23. Delaney J, Hodson MP, Thakkar H, Connor SC, Sweatman BC, Kenny SP, McGill PJ, Holder JC, Hutton KA, Haselden JN, Waterfield CJ. Tryptophan-nad+ pathway metabolites as putative biomarkers and predictors of peroxisome proliferation. Arch Toxicol. 2005;79:208–223

24. May DH, Navarro SL, Ruczinski I, Hogan J, Ogata Y, Schwarz Y, Levy L, Holzman T, McIntosh MW, Lampe JW. Metabolomic profiling of urine: Response to a randomised, controlled feeding study of select fruits and vegetables, and application to an observational study. Br J Nutr. 2013;110:1760–1770

25. Lloyd AJ, Beckmann M, Fave G, Mathers JC, Draper J. Proline betaine and its biotransformation products in fasting urine samples are potential biomarkers of habitual citrus fruit consumption. Br J Nutr. 2011;106:812–824

26. Aronov PA, Luo FJ, Plummer NS, Quan Z, Holmes S, Hostetter TH, Meyer TW. Colonic contribution to uremic solutes. J Am Soc Nephrol. 2011;22:1769–1776

27. Wang Y, Lawler D, Larson B, Ramadan Z, Kochhar S, Holmes E, Nicholson JK. Metabonomic investigations of aging and caloric restriction in a life-long dog study. J Proteome Res. 2007;6:1846–1854

28. Lennard L, Lilleyman JS, Van Loon J, Weinshilboum RM. Genetic variation in response to 6-mercaptopurine for childhood acute lymphoblastic leukaemia. Lancet. 1990;336:225–229

29. Kirchheiner J, Nickchen K, Bauer M, Wong ML, Licinio J, Roots I, Brockmoller J. Pharmacogenetics of antidepressants and antipsychotics: The contribution of allelic variations to the phenotype of drug response. Mol Psychiatry. 2004;9:442–473

30. Beynen AC, Katan MB, Van Zutphen LF. Hypo- and hyperresponders: Individual differences in the response of serum cholesterol concentration to changes in diet. Adv Lipid Res. 1987;22:115–171

31. Stamler J, Elliott P, Appel L, Chan Q, Buzzard M, Dennis B, Dyer AR, Elmer P, Greenland P, Jones D, Kesteloot H, Kuller L, Labarthe D, Liu K, Moag-Stahlberg A, Nichaman M, Okayama A, Okuda N, Robertson C, Rodriguez B, Stevens M, Ueshima H, Horn LV, Zhou B. Higher blood pressure in middle-aged american adults with less education-role of multiple dietary factors: The intermap study. J Hum Hypertens. 2003;17:655–775

